# Machined silicon traps for capturing novel bacterial communities and strains *in-situ*

**DOI:** 10.1101/2023.02.21.529475

**Authors:** Clara Romero Santiveri, Joseph H. Vineis, Sofia Martins, Carlos Calaza, João Gaspar, Jennifer L. Bowen, Edgar D. Goluch

## Abstract

We tested the feasibility of a novel machined silicon nanopore enrichment device to recover individual microbial taxa from anaerobic sediments. Unlike other environmental isolation devices that have multiple entry points for bacteria or require the sample to be manually placed inside of a culturing chamber, our silicon device contains 24 precisely sized and spaced nanopores, each of which is connected to one culturing well, thereby providing only one entry point for bacteria. The culturing wells allow nutrient transport, so the bacteria that enter continue to experience their natural chemical environment, allowing collection of microbes without manipulating the environment. The device was deployed in marsh sediment and subsequently returned to the laboratory for bacterial culturing and analysis. 16S rRNA marker gene and metagenomic sequencing was used to quantify the number of different microbial taxa cultured from the device. The 16S rRNA sequencing results indicate that each well of the device contained between 1 and 62 different organisms from several taxonomic groups, including likely novel taxa. We also sequenced the metagenome from 8 of the 24 wells, enabling the reconstruction of 56 metagenomic assembled genomes (MAGs), and 44 of these MAGs represented non-redundant genome reconstructions. These results demonstrate that our novel silicon nanofluidic device can be used for isolating and culturing consortia containing a small number of microbial taxa from anaerobic sediments, which can be very valuable in determining their physiological potential.

**Importance:** There are very few methods that can remove a few bacterial cells from a complex environment and keep the cells alive so that they can propagate sufficiently to be analyzed in a laboratory. Such methods are important to develop because the physiological functions of individual species of bacteria are often unknown, cannot be determined directly in the complex sample, and many bacterial cells cannot be grown outside of their natural environment. A novel bacterial isolation device has been made tested in a salt marsh. The results show that the device successfully isolated small groups of bacterial species from the incredibly diverse surroundings. The communities of bacteria were easily removed from the device in the laboratory and analyzed.

## Introduction

Microorganisms are the most diverse forms of life on Earth. There are 100 million times as many bacterial cells in the oceans (13×10^28^) as stars in the known universe [1]. Despite the astonishing progress in microbiology over the past century, we have only scratched the surface of this enormous microbial world. It has been estimated that *<*1% of bacterial species have been cultured in the laboratory [2]. Based on sampling location, *∼*1% of sediment bacteria, 0.01-0.1% of soil bacteria, and 0.001-0.1% of marine (surface) bacteria have been cultivated in the laboratory [3]. To improve and accelerate bacterial cultivation, microfluidic devices with various configurations have been developed for sorting, isolating, and studying microorganisms (Table 1). However, most microfluidic devices require sophisticated external instrumentation to be operated (active microfluidics) and, therefore, need to remove the sample from the original environment for processing, which potentially introduces sample bias and loss of diversity [4-8]. These active microfluidic techniques manipulate the particles’ movement in real-time by using external forces, including electric fields [9-12], acoustic streaming [13], magnetic fields [14-15].

**Table 1.** Summary of articles that used different microfluidic-based approaches for cell sorting and isolation..

| Isolation process | References | Microfluidic type | Does it disturb the environment? |
| --- | --- | --- | --- |
| Droplet-based | Watterson *et al.* (2020) [4] | Active | Yes |
|  | Eun *et al.* (2011) [5] |  |  |
|  | Villa *et al.* (2019) [6] |  |  |
|  | Leung *et al.* (2012) [7] |  |  |
| Pressure-driven | Bamford *et al.* (2017) [8] | Active | Yes |
| Dielectrophoresis | Lu *et al.* (2013) [9] | Active | Yes |
|  | Jiang *et al.* (2019) [10] |  |  |
|  | D'Amico *et al.* (2017) [11] |  |  |
|  | Braff *et al.* (2013) [12] |  |  |
| Acoustophoresis | Dow *et al.* (2018) [13] | Active | Yes |
| Magnetic beads | Chang *et al.* (2014) [14] | Active | Yes |
|  | Miller *et al.* (2019) [15] |  |  |
| Dilution | Nichols *et al.* (2010) [16] | Active | Yes |
|  | Yoshiteru *et al.* (2009) [17] |  |  |
|  | Berdy *et al.* (2017) [18] |  |  |
| Microfiltration | Raub *et al.* (2015) [19] | Passive | Yes |
|  | Fan *et al.* (2015) [20] |  |  |
| Selective lysis | Zelenin *et al.* (2015) [21] | Passive | Yes |
| Inertial (flows) | Wu *et al.* (2009) [22] | Passive | Yes |
| Chemotaxis | Männik *et al.* (2009) [23] | Passive | Yes |
|  | Tandogan *et al.* (2014) [24] |  | No |
|  | Rezaei *et al.* (2019) [25] |  | No |
|  | Our device |  | No |

A few passive sorting microfluidic devices have been demonstrated, but only a few do not disturb the environment. The iChip, for example, has been successfully used to cultivate many new species of bacteria, however, the sample must be collected by the user, diluted, the cells then placed inside of the isolation chambers, prior to placing it into the environment for nutrient exchange [16-18]. Completing these steps in the field is cumbersome, and placement of the device back in exact location, to the millimeter, where the sample was collected is nearly impossible. Rezaei *et al*. designed an ingestible pill device recently, which does not disturb the environment as it takes samples from the gut after being swallowed [25]. However, the purpose of the device is different. The ingestible pill is intended for sampling of gut microbiota and it does not limit the bacterial diversity that is collected.

Tandogan *et al*. developed a polymer nanofluidic device, a predecessor of the device demonstrated in this article, which used a similar design with sub-micrometer channel features to limit bacterial cell access isolation chambers [24]. Our device overcomes several limitations from the previous version. Anaerobic bacteria can now be cultured using silicon wafers and polycarbonate as the central part of the trap instead of polydimethylsiloxane (PDMS), which is gas permeable. The constrictions in our device are exposed directly to the environment sample without the bacteria needing to enter the main channel before getting to the constrictions, whereas the PDMS device required that the bacterial cells travel nearly a centimeter to reach the nanochannel. Finally, and most importantly, the silicon nanopore devices can be manufactured in volume using established microfabrication techniques borrowed from the microelectronics industry.

Here, we describe a novel, passive, nanofabricated device that allows for in-situ isolation of bacterial species. The isolated bacteria are exposed to nutrients in their natural surroundings using a nanoporous membrane. Thus, the device eliminates the need for sample processing before initiating a culture and provides the opportunity to perform genomic analysis on cells obtained directly from natural communities.

A silicon single-side polished (SSP) wafer and a silicon on insulator (SOI) wafer are used as the base of the device; it has 24 holes (constrictions) that vary in diameter, they range from 2 µm to 0.5 µm on the SSP and from 1.1 µm to 0.1 µm on the SOI wafer. These constrictions are at least one dimension smaller than the diameter of a bacterial cell. Fresh food in the isolation chambers chemotactically attracts microorganisms toward the constrictions (Figure 1). As a result, bacterial species get trapped at the entrance of these sub-micron constrictions (Figure 1B), preventing other bacterial cells from reaching the isolation chamber. The trapped microorganism continues to divide (Figure 1C), and each progeny advances further through the constriction. Finally, after several successions, only one species will enter the isolation chamber, which is the predecessor of the trapped species (Figure 1D).

**Figure 1.**
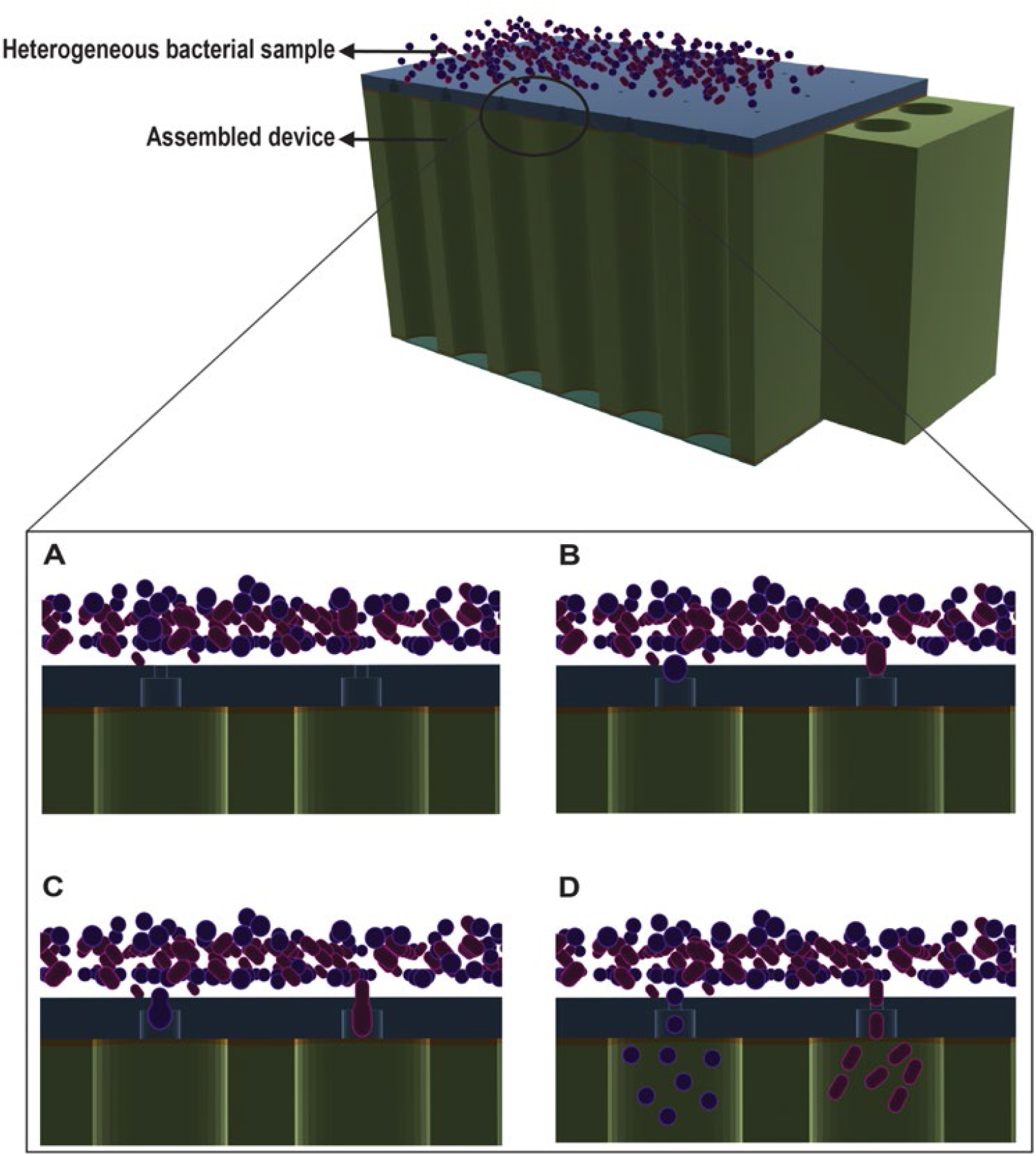
Schematic of the microfluidic device. The wafer is connected to the isolation chambers via nano- and sub-micron constrictions. Heterogeneous bacterial culture self-sort into different isolation chambers with the help of chemotaxis and size-specific constrictions.

Microbial diversity and community composition is assessed using 16S ribosomal RNA (rRNA) sequencing. This technique has allowed the discovery of important relationships between microbial structure and function and led to the discovery of the “rare biosphere” [26]. However, estimates of diversity and species counts can be heavily influenced by the differences in the number of 16S rRNA operons within individual organisms [27], polymerase chain reaction (PCR) errors, sequencing errors [28], and primer bias [29]. Genome reconstruction from metagenomic data can provide a less biased representation of the diversity of a community because the preparation of samples requires fewer PCR cycles, avoids primer bias, and analytical approaches are not influenced by the operon structure of individual marker genes. This approach also allows us to identify the metabolic potential of microbial organisms within the environment. However, the immense diversity of natural communities hampers our ability to reconstruct all microbial genomes from most environmental samples.

While dilution to extinction and enrichment cultures are commonly used to overcome this problem, they are generally conducted within the laboratory under purely synthetic conditions. The ability to isolate a reduced community or individual strains *in-situ* significantly increases the opportunity for microbiologists to identify novel microbial metabolism and interactions. *In-situ* isolation can also improve current laboratory cultivation yield because the metabolic handoffs and environmental conditions relied upon by many taxa for growth are preserved. Further, pure bacterial cultures are essential for understanding investigating virulence factors, antibiotic susceptibility, and genome sequences. However, only a few bacterial species can be cultivated by routine culture, so molecular analyses of environmental sequences are employed to substantially expand our knowledge of microbial life [30], [31].

## Materials and Methods

### Wafer Fabrication

The micromachining of silicon-on-insulator (SOI) wafer substrates was performed using a SiO2 hard mask for the silicon dry etching process. A 1.5 µm thick SiO2 layer was first deposited on the front side of the SOI substrate using a plasma-enhanced chemical vapor deposition (PECVD) system (MPX from SPTS). Direct write laser lithography (DWL 2000 from Heidelberg Instruments) with a 1.2 µm thick AZP4110 positive photoresist was then used to define the geometry of the small constrictions on the front side. The layout consisted of a 6×4 matrix of circular constrictions with diameters ranging from 1.50 µm to 2.25 µm. The layout was repeated 28 times on an 8-inch-diameter wafer. After exposure, the resist was developed using AZ400K. The SiO_2_ layer was then patterned by reactive ion etching (RIE) using a C_4_F_8_ based plasma in an APS reactor from SPTS. Next, 10-µm-deep circular constrictions were achieved using deep reactive ion etching (DRIE) of silicon (Pegasus system from SPTS) using a SF_6_/C_4_F_8_ etching chemistry. The resist mask was then stripped using an oxygen plasma etch (PVA GIGAbatch 360 M tool from Tepla).

The backside of the wafer was micromachined as well to achieve through-wafer channels. First, a 5-µm-thick PECVD SiO_2_ layer was deposited on the back of the wafer. This layer acted as a hard mask for the subsequent DRIE step. Next, lithography for the patterning of through-wafer channels was performed on a Mask Aligner system (MABA6 from SussMicroTec) using a 2.2 µm thick AZP4110 positive photoresist. The layout consisted of a 6×4 matrix of circular holes with a diameter of 25 µm, aligned with the previous frontside lithography and repeated 28 times. After exposure, the resist was developed using AZ400K.

The processes used for both the etching of the SiO_2_ hard mask and the resist strip are similar to those previously used on the front side. Before the DRIE process, a thermal release tape was applied on the wafer’s front side to prevent leakage through the front side constrictions in the event of breakage on the SOI buried oxide layer (BOX). The through-wafer channels with 725 µm depth were obtained with a DRIE process using a SF_6_/C_4_F_8_, which stopped on the BOX layer. Then the thermal release tape was removed by placing the wafer on a hotplate at 180 °C, and channels were opened by removing the exposed BOX layer using an HF vapor tool (Primaxx from SPTS).

The size of the small constrictions can be tailored for different applications. For example, if narrower constrictions are required, a new PECVD SiO_2_ layer can be deposited to reduce the effective diameter of the SiO_2_. A 2.1-µm-thick layer was deposited in this case to obtain constrictions with dimensions in the range of 0.25 µm - 1.25 µm.

Dicing the wafers into 28 individual devices containing the 6×4 array of channels without damaging the small constriction structures was a uniquely challenging step. This process was completed by assembling a protection thermal release tape on the front side and a regular dicing tape on the backside of the wafer and performing the dicing from the front side. Dicing tape was then released with UV exposure for a few minutes and front side tape using heating the wafer in an oven at 180 °C. A process diagram for device fabrication is provided in the Supporting Information as Figure S1.

A silicon single-side polished (SSP) wafer and a silicon on insulator (SOI) wafer are used as the base of the device; it has 24 holes (constrictions) that vary in diameter, they range from 2µm to 0.5µm on the SSP and from 1.1µm to 0.1µm on the SOI wafer.

The final wafer can be divided into rows; there are four rows with six constrictions on each row (Figure 2) ad each row has different constriction diameter (Table 2). The SSP wafer constriction diameters range from 2.0 µm to 0.5 µm while the SOI wafer diameters range from 1.0 µm to 0.1 µm.

**Figure 2.**
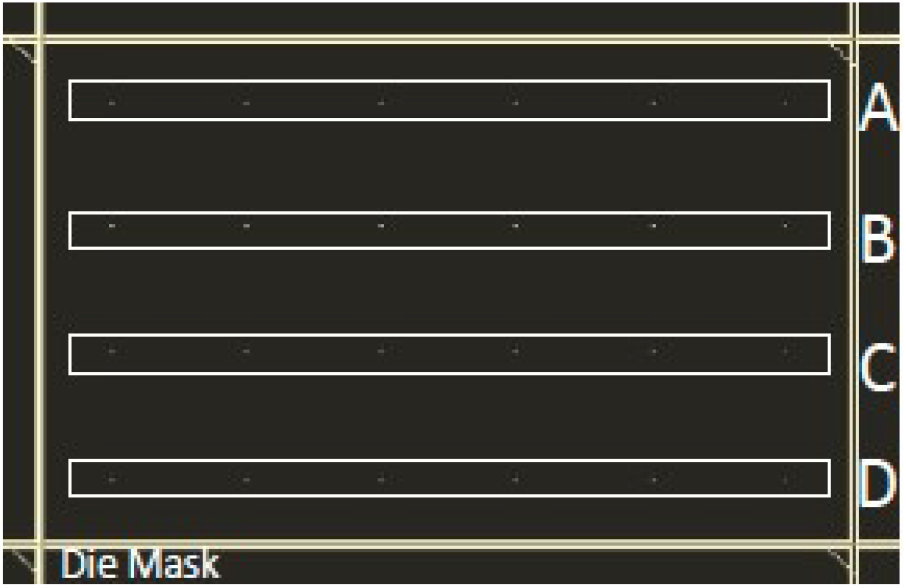
Arrangement of constrictions on a silicon device.

**Table 2.** Constriction diameters.

| Wafer | A | B | C | D |
| --- | --- | --- | --- | --- |
| SSP | 2$\mu\text{m}$ | 1.4$\mu\text{m}$ | 1$\mu\text{m}$ | 0.5$\mu\text{m}$ |
| SOI | 1.1$\mu\text{m}$ | 0.7$\mu\text{m}$ | 0.4$\mu\text{m}$ | 0.1$\mu\text{m}$ |

### Device Assembly

The device consists of 4 elements (Figure 3): the wafer, two double-sided adhesives (Adhesive Transfer Tape Acrylic Adhesive Clear, DigiKey), a polycarbonate body (Clear Impact-Resistant Polycarbonate, McMaster-Carr), and a Nuclepore track-etched polycarbonate (PC) membrane (0.05 µm pore size, Whatman). Before assembling the final device, the adhesive was cut using an Epilog Zing laser cutter (30W). Next, circles were cut in the double-sided adhesive so that there was an open path between the channels in the wafer and the wells in the polycarbonate and between the wells and the nanoporous membrane. Finally, the nanoporous membrane was manually cut to match the size of the polycarbonate part.

**Figure 3.**
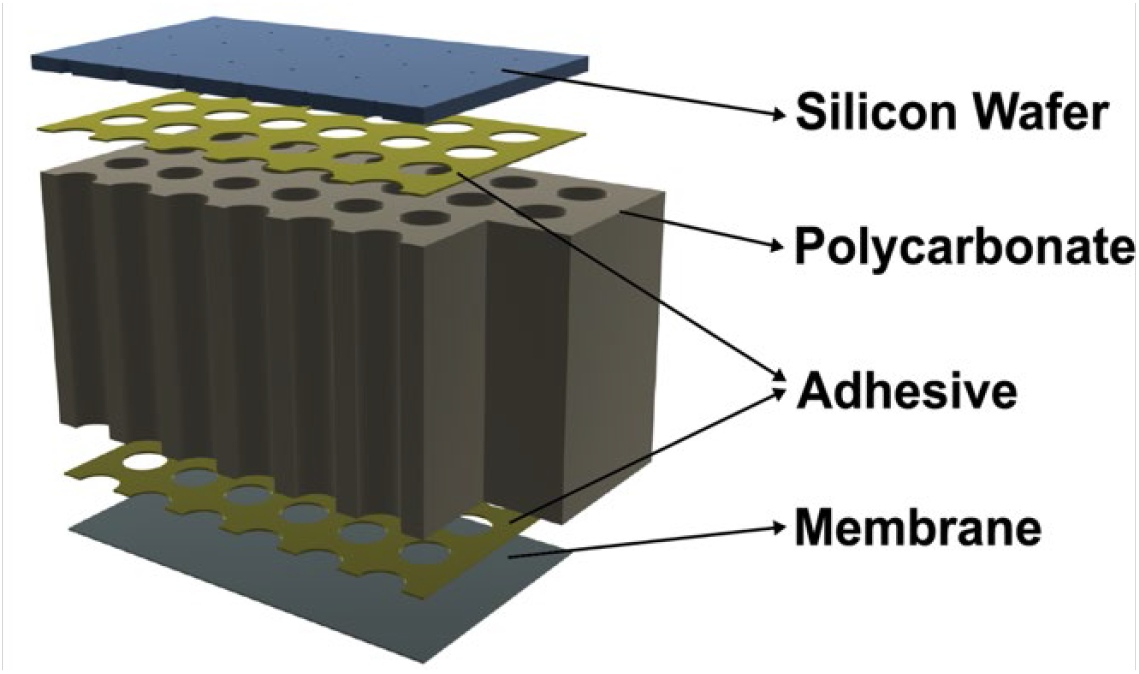
Schematic identifying the parts off the silicon trap.

After autoclaving all of the components, the double-sided adhesive was first adhered to the polycarbonate body aligning the holes and wells. Then, one side of the polycarbonate, the rough backside of the wafer, was attached to the polycarbonate, while the smooth front side was exposed to the environment. Finally, the wafer and the plastic part were designed to align correctly using hand positioning.

Next, the device was filled with liquid medium, and finally, the nanoporous membrane was attached to the polycarbonate body using a second piece of double-sided adhesive. The membrane pore size is small enough to block bacterial cells from entering the isolation well but wide enough to allow diffusion of nutrients into the device to cultivate the trapped bacteria.

### Cultivar Collection

A single trap consisting of 24 chambers, basic marine medium (Gibco), a polycarbonate membrane, and the silicon wafer were autoclaved, sterilized, and placed into an anaerobic chamber (Coy) containing 95% N_2_ and 5% H_2_ gas in the presence of a Stak-Pak catalyst for 48 hrs to remove O_2_.

Nitrate was added to the medium after sterilization to a final concentration of 1000 µM. After attaching the bottom of the trap with adhesive, we added approximately 50 µL of medium to each well before adding the nutrient permeable membrane on the top. The trap was placed into a 50 mL conical tube filled with the same medium and transported to Plum Island Long Term Ecological Research (LTER) Site, approximately 1 hr away from the Northeastern University Marine Science Center. A soil core (8 × 40 cm) was taken from the sediment of the short ecotype of *Spartina alterniflora* on the high marsh platform. A sterile razor blade was used to make an incision approximately 4 cm deep along the core length. The device was embedded into the core at 35 cm from the surface, and the entire core was returned to its original position and allowed to incubate for 10 days. After the incubation period, the traps and surrounding sediment were recovered, placed into a plastic bag, and transferred to an anaerobic chamber within an hour. The trap was rinsed with sterile deionized water and cleaned with 70% ethanol using a Kimwipe. A sterile 1 ml syringe was used to transfer the entire contents of each well to separate Hungate tubes containing 10 mL of sterile basic marine medium. Several Hungate tubes containing medium were not inoculated to serve as negative controls. Growth was determined by turbidity and the presence of black particulates in the medium that likely resulted from sulfur-driven iron reduction.

### DNA purification

After 21 days of growth in the Hungate tubes, we purified DNA from 1 mL of cells and medium using a sucrose lysis buffer approach adapted from Britschgi and Fallon 1994 [32]. In addition, duplicate DNA extractions were completed for four of the samples to assess extraction and PCR bias.

### 16S amplification and ASV clustering

Partial 16S rRNA gene sequences were amplified from the purified DNA according to Caporaso et al. [33] and sequenced on a MiSeq using 2 × 250 PE v2 chemistry. Reads were quality filtered, merged, and clustered into amplicon sequence variants (ASVs) using the Dada2 pipeline v 1.14.0.[39].

### Metagenomic library construction and MAG reconstruction

Metagenomic libraries were constructed for eight of the cultures that displayed unique combinations of ASVs. We sheared approximately 1 µg of purified DNA as input for the NuGen Ovation R DNA library prep kit and followed the recommendations of the protocol to create all libraries. Each library was quantified using the Invitrogen pico-green DNA assay, and we pooled all eight libraries based on the picogreen concentrations in an equimolar fashion. We size selected the pooled libraries to 600 bp using a Covaris ME220 ultrasonicator according to the manufacturer’s recommendations. The library was cleaned using AMPure XP R DNA purification beads at a 1:1 DNA to bead ratio. We quantified the final library using a Kapa qPCR Illumina library quantification kit to optimize the concentration of the library for sequencing. The library was sequenced on an Illumina MiSeq according to PE 2 × 250 v3 chemistry. All reads were quality filtered using Illumina-utilities v2.6 using the default parameters of “iu-filter-minoche” [35]. Filtered reads were assembled into contigs using the SPAdes genome assembler v3.13.0 [36] according to the metagenomic pipeline. Finally, we mapped the short reads from each of the eight samples onto each of the individual assemblies using bowtie2 v 2.2.9 [37].

We used Anvi’o v 6.1 [38] to reconstruct genomes from the assembled metagenomic data. We began by creating a contigs database using the command “anvi-gen-contigs-database,” which included identification of open reading frames (ORFs) using Prodigal [39], calculation of contig tetranucleotide frequency, and splitting contigs larger than 20 kbp into 10 kbp “splits.” The command “anvi-run-hmms” searched all contigs for the presence of single-copy genes using three separate collections, including bacterial, archaeal, and eukaryotic collections. This algorithm uses HMMER as the search engine to identify the presence of single-copy gene collections [40]. To link the mapping data for each sample to the contigs database, we used the command “anvi-profile.” All profile databases were merged using “anvi-merge,” and we used a manual approach employed by “anvi-interactive” to place contigs into bins that were most similar in coverage profiles across all samples.

The interactive interface of Anvi’o also allowed us to evaluate the percentage of single-copy genes detected and those that were redundant in the collection of contigs to more accurately place contigs into MAGs. Filtered sequencing reads are contained within NCBI under the project PRJNA714626. The specifications of each command can be found here (https://github.com/jvineis/Enrichment-Traps), and the files required to visualize the selection of contigs can be found here (https://doi.org/10.6084/m9.figshare.13650800). We created a list of non-redundant MAGs based on their average nucleotide identity (ANI) using two steps. First, we ran “anvi-compute-genome-similarity” to calculate the pairwise percent identity and the percent alignment of all MAGs. Then we used “anvi-dereplicategenomes” to identify MAGs that contained 95% ANI across 90% of their genome, specifying the use of pyANI [41]. Finally, we identified MAG taxonomy using “anvi-run-scg-taxonomy,” which uses DIAMOND [42] to search single-copy genes identified in the MAGs to reference sequences in the Genome Taxonomy Database (GTDB) [43].

### Estimating MAG relative abundance

Following MAG reconstruction and dereplication, we exported a fasta file for each split in the collection of MAGs and mapped each of the short read metagenomic datasets back to this fasta file using bowtie2. We converted the resulting sam file to a bam file and removed all alignments with a MAPQ score below 10 using “samtools view.” Removal of alignments below this threshold is an effective way to remove non-specific alignments and reads that map to more than one position. However, multiple alignments for individual reads can still be retained using this method which can slightly influence relative abundance estimates. We tabulated the number of reads that were recruited to each split using “samtools idxstats” and a custom script to tabulate the number of reads for each MAG.

## Results and Discussion

The 16S rRNA sequencing effort produced an average of 22,879 high-quality reads per sample with a minimum of 14,513 and a maximum of 28,780 (Figure 4). A total of 185 unique ASVs were detected, and the number of ASVs per sample ranged from 1 to 62, with a mean of 23 (Figure S2). The technical replicate amplicon processing from four samples (indicated by colored boxes at the bottom of Figure 4) indicates that the results are robust for separate DNA extractions of the same culture.

**Figure 4.**
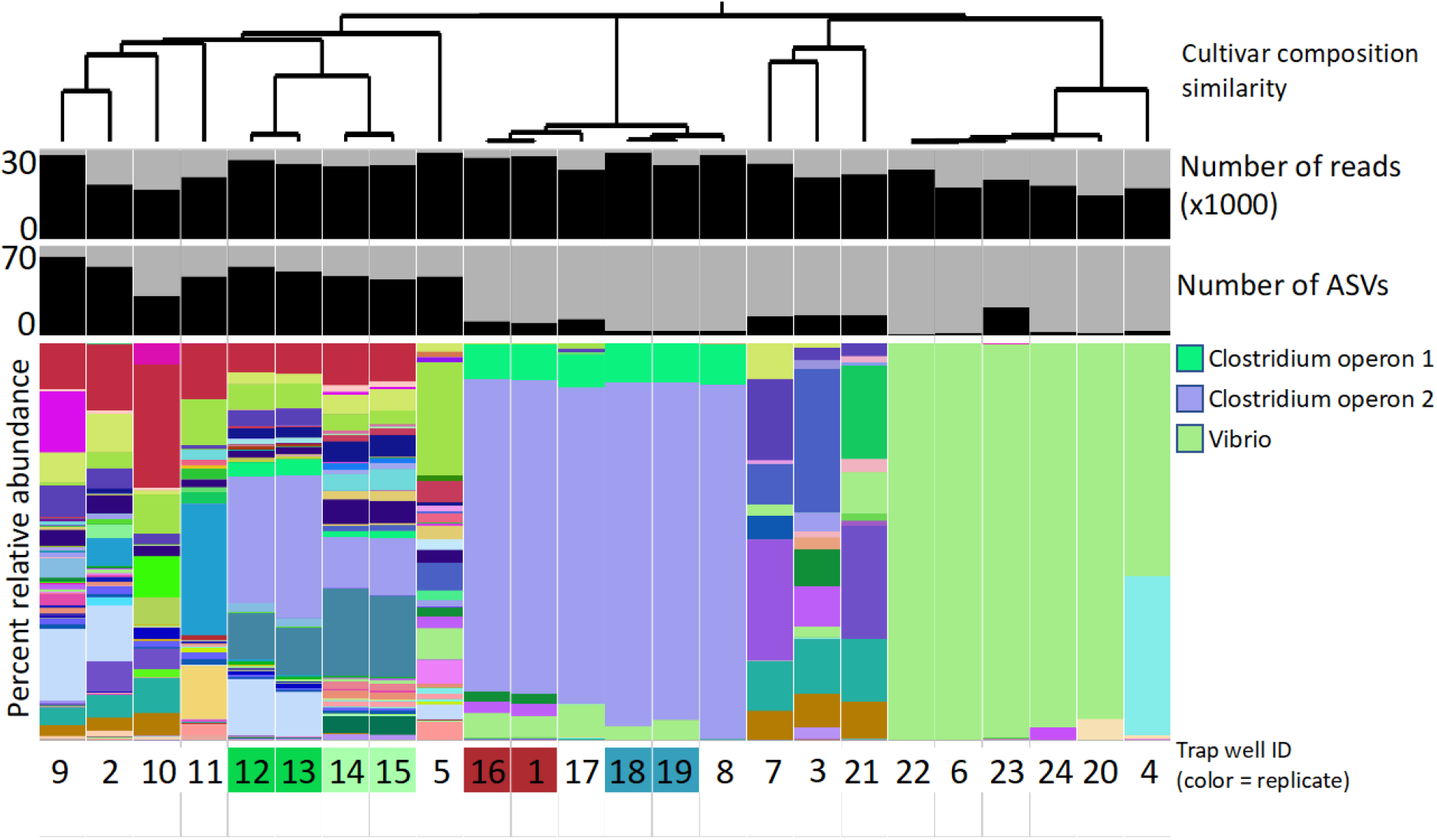
Summary of ASVs detected in trap wells, including a hierarchical tree showing the relationship in ASV composition among each of the trap wells (top). Barplots include the number of quality filtered reads per sample (top) the number of ASVs (middle), and the percent relative abundance of each ASV within each of the trap wells (bottom). The number at the bottom of the figure indicates the ID of the well. The presence of color behind the number indicates if the sample was a technical replicate and if two boxes have the same color then the DNA was extracted from the same cultivar.

The diversity of ASVs within the wells of the trap can be broken down into three major groups. In the first group (trap well numbers 22, 6, 23, 24, and 20), a single ASV most closely related to *Vibrionaceae* represented more than 94% of all sequences (Figure 2). In two of the trap wells (22 and 6), this ASV represented more than 99% of all sequences. The second group, representing four of the trap wells (17, 18, 19, and 8), is dominated by two ASVs that are likely operons derived from the same organism with taxonomic resolution to *Clostridiales*. An alignment of the two representative sequences for the *Clostridiales* ASVs indicated that there was a single nucleotide difference between them, and they occurred at a 9:1 ratio within all samples where they were detected. The two ASVs combined to reach greater than 85% of all sequences in four of the trap-wells. The third group was comprised of trap wells containing a diversity of bacterial taxa. Within this group, there were 127 ASVs that occurred in less than three samples, and 87 of these were never detected above 5% in any of the trap wells (Figure S2, Table S1). The remaining 55 ASVs occurred in three or more samples, and the mean percent relative abundance for this group of ASVs was 3.8. Twelve ASVs that occurred in more than two wells had a mean of 5 percent relative abundance (Figure 4, Figure S2, Table S1). These results indicate that there was significant overlap in the cultured organisms isolated from many of the trap wells, which is surprising given the large amount of diversity that exists within salt marsh sediments [44], [45]. We reconstructed a total of 56 draft genomes of medium to high quality according to MIMAG standards [46] from eight of the trap wells. Dereplication of these MAGs produced a set of 44 unique MAGs (Table S2).

In trap well #8, where ASV analysis indicated the presence of a dominant organism closely related to *Vibrio*, a single MAG was resolved with completion and redundancy scores of 100% and 0%, respectively, with 68% of all short reads recruiting back to the MAG and consistent coverage across all contigs with the exception of a contig containing 16s rRNA genes (Figure 5). Two additional MAGs with 95% ANI across 90% of their genome were recovered from two other wells and likely represented organisms from the same population (Figure 5). In trap wells with a greater diversity of organisms, we recovered up to 17 MAGs with over 50% read recruitment in nearly all samples (Figure 5).

**Figure 5.**
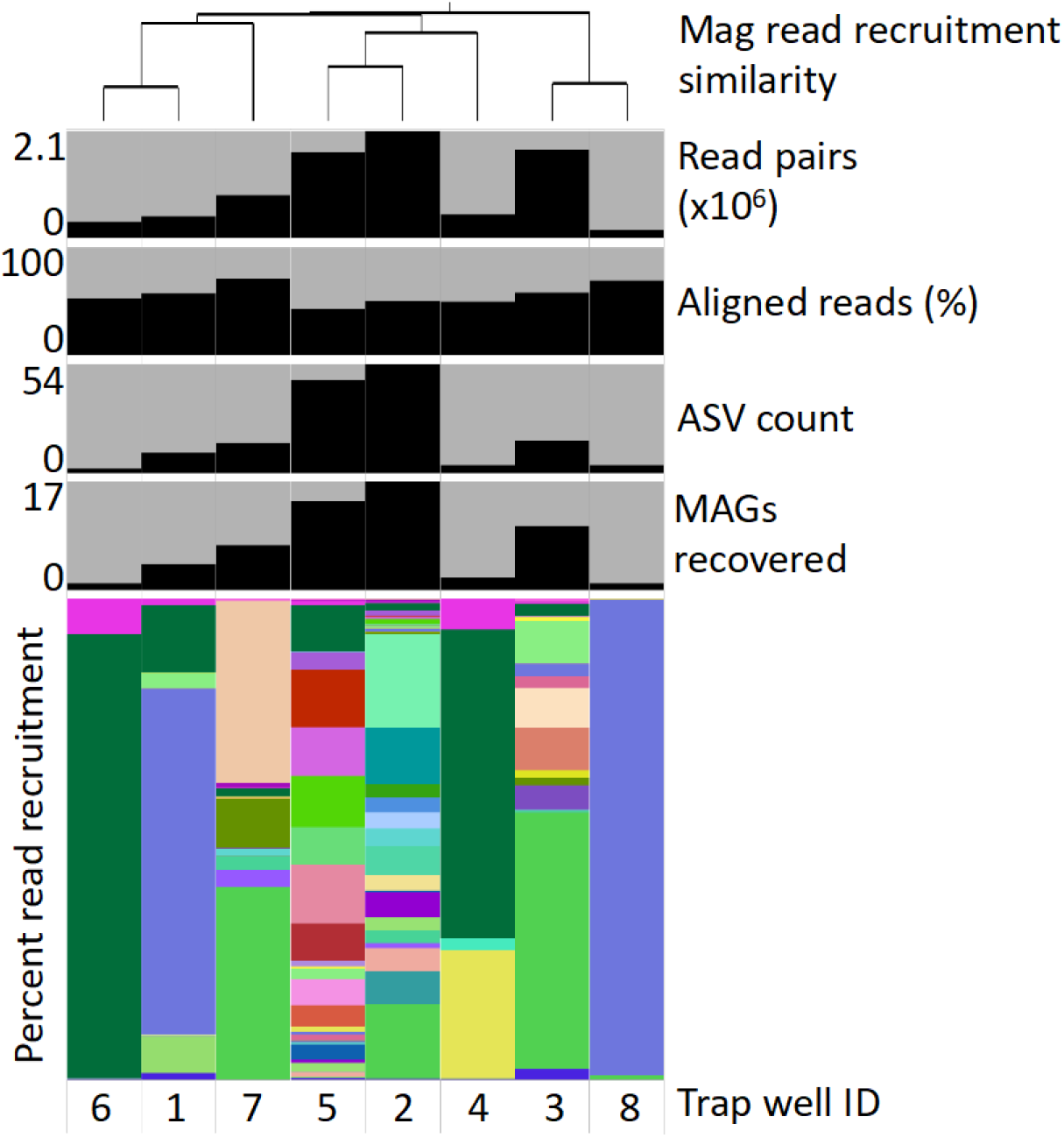
Summary of metagenomic assembled genomes (MAGs) from eight trap wells ordered by the hierarchical tree (top) based on the similarity in relative percent read recruitment of all MAGs. Bar plots, from top to bottom indicate 1) the number of quality filtered read pairs per sample, 2) the number of reads that align to the non-redundant collection of MAGs, 3) the number of ASVs detected in the sample, 4) the number of MAGs recovered from each sample, and 5) the percent relative abundance of each non-redundant MAG in each of the cultivars. The trap wells are identified at the bottom of the figure.

We observed minimal variability in alignment of short reads too many of the MAGs in this study (Figure 6), indicating that in most cases, we were able to isolate individual strains.

**Figure 6.**
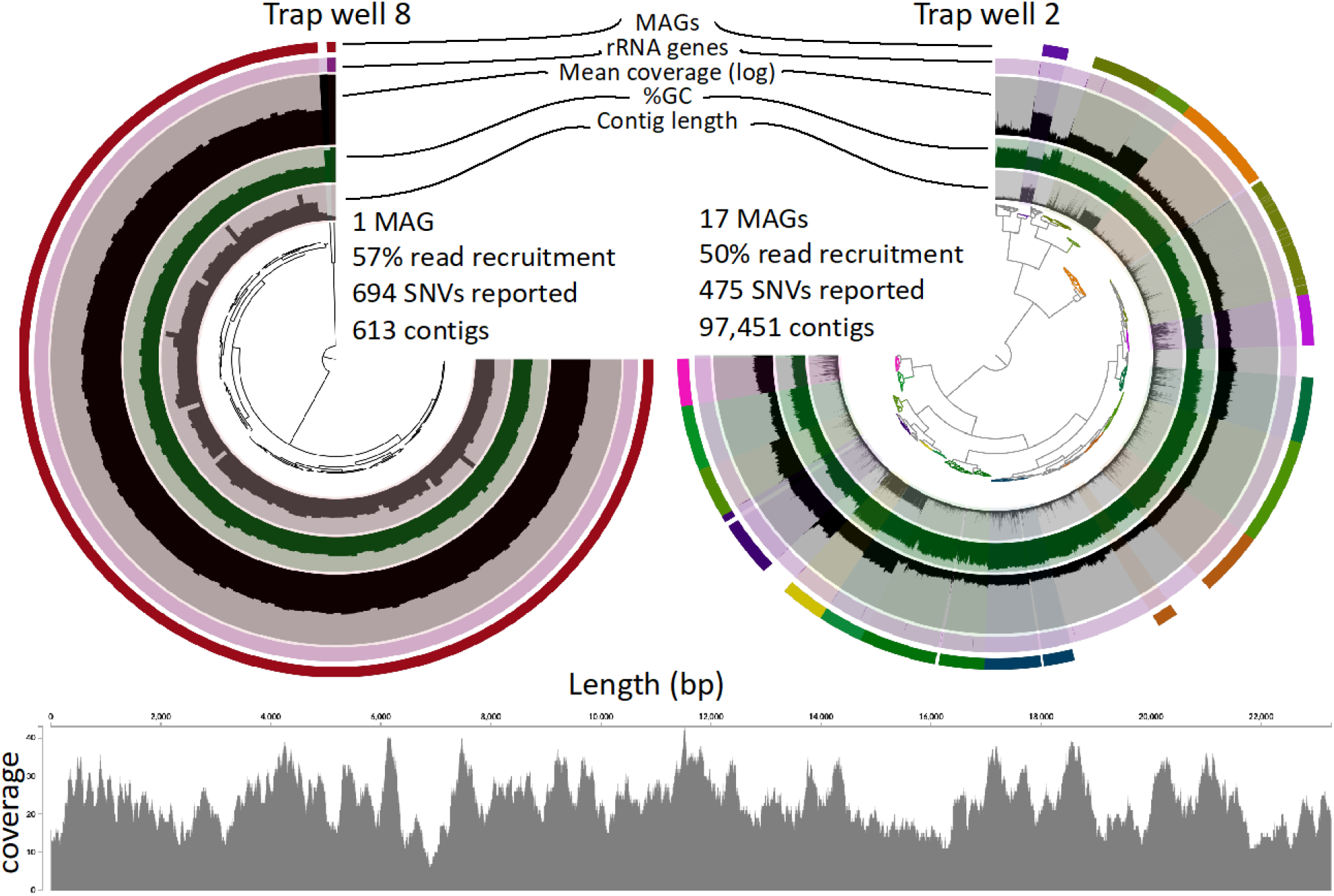
Comparison of MAGs from two traps. Each circular display contains a tree at the center representing the similarity of tetranucleotide frequency and coverage of each contig in the two independent assemblies. Subsequent layers demonstrate 1) contig length, 2) % GC content, 3) log mean coverage of the contig within the sample, 4) an indicator of whether a 16s rRNA gene was detected in the sample and 5) the bin location of the MAG collection. The coverage profile (bottom) shows an example of one 22 kbp contig derived from one of the MAGs. Any variation in the consensus of short reads mapping back to this contig would be highlighted and absence of any variation indicates that the short reads completely agree with the consensus.

The number of MAGs identified within the traps was highly correlated with the number of ASVs detected, and the relationship between the number of ASVs and MAGs was linear, with nearly three times the number of ASVs observed for every MAG (Figure 7).

**Figure 7.**
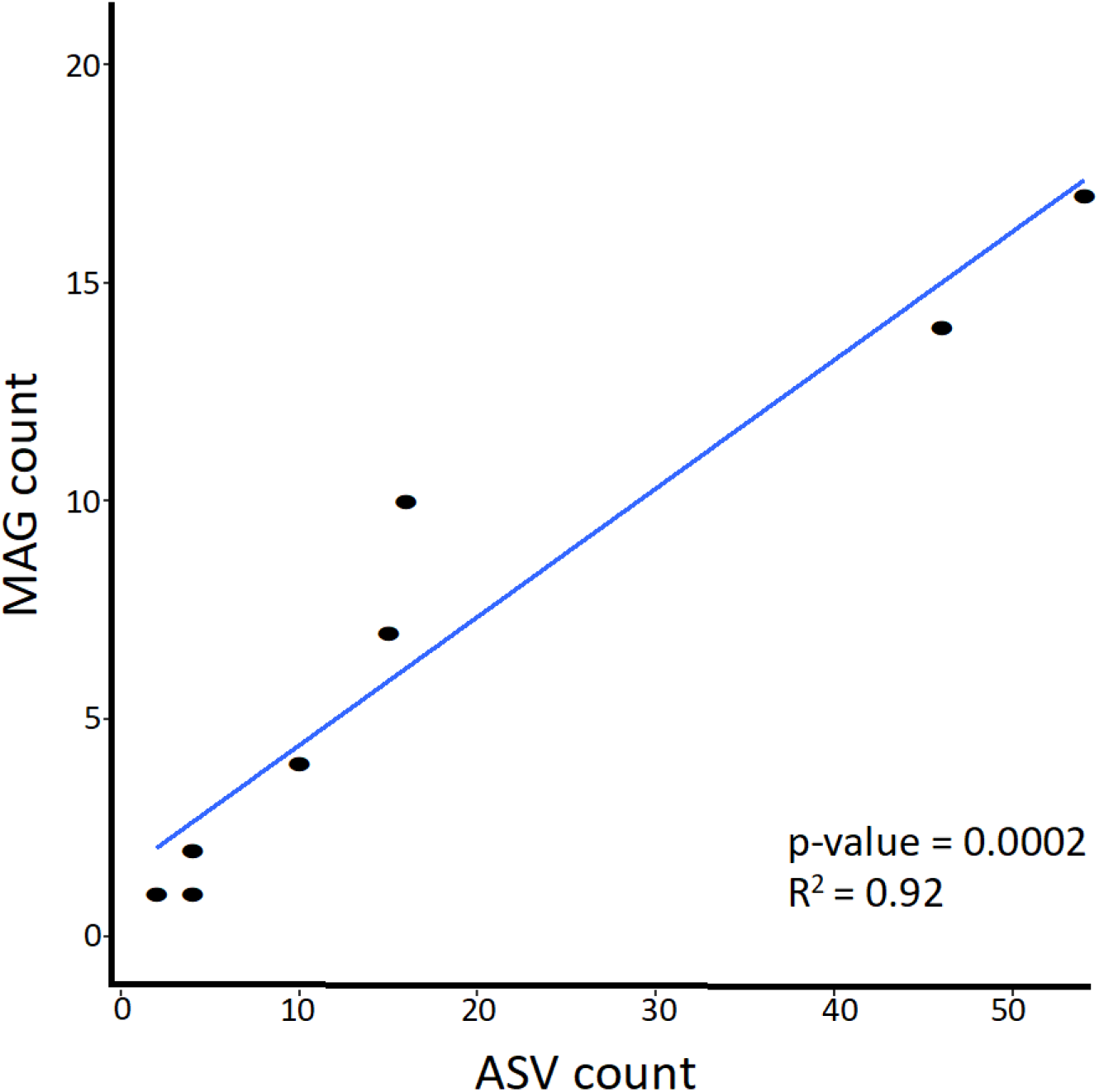
Linear model describing the relationship between the number of ASVs and MAGs recovered from each trap well.

This could result from the presence of multiple operons of the 16S rRNA gene within several of the genomes. This result indicates that 16S rRNA amplicon sequencing may overestimate the number of organisms isolated in each well, and estimates can be improved by metagenomic sequencing and genome reconstruction. We recovered ASVs and MAGs that could not be assigned taxonomy to the family level, indicating that they represent novel organisms. Obtaining genomic information for these organisms is a significant step toward understanding their functional capacity and provides us with the culture collections to validate their physiological potential. This system offers a considerable improvement to classical approaches of dilution to extinction and streaking plates because it allows for the capture of communities and strains in-situ with the potential to use multiple media types in the same trap.

## Acknowledgment

This material is based upon work supported by the National Science Foundation under grant no. IDBR 1353853. Support for the amplicon and metagenomic sequencing was provided by NSF grant no. DEB 1350491 to JLB. PIE LTER, where these samples were collected, is supported by NSF Grant no. OCE 1637630. EDG has a financial interest in the trap technology. Traps can be obtained by researchers through Microbial Devices, LLC.

